# *Drosophila melanogaster* model of RVCL-S demonstrates age dependent disease progression

**DOI:** 10.1101/2025.11.10.687584

**Authors:** Elena Gracheva, Abigail Matt, Fei Wang, Raymond Hsin, Hongwu Liang, Xiangping Ouyang, Jimin Ding, Jonathan J. Miner, Chao Zhou

**Affiliations:** Department of Biomedical Engineering, Washington University in St Louis, 1 Brookings Dr, St Louis, MO 63130, USA; Department of Statistics and Data Science, Division of Biostatistics, Washington University in St Louis, 1 Brookings Dr, St Louis, MO 63130, USA; Departments of Medicine and Microbiology, RVCL Research Center, and Colton Center for Autoimmunity, University of Pennsylvania Perelman School of Medicine, 522B Johnson Pavilion, 3610 Hamilton Walk, Philadelphia, PA 19104 USA

**Keywords:** Drosophila melanogaster, Optical coherence microscopy (OCM), RVCL-S, RVCL, TREX1, cg3165

## Abstract

Retinal vasculopathy with cerebral leukoencephalopathy and systemic manifestations (RVCL-S) is a disease that causes deterioration of small vessels, affecting various organs: eyes, brain, liver, and others. The RVCL-S carriers have lower life expectancy. There is no cure available to date. The disease has been linked to mutations in TREX1 gene disrupting its cytoplasmic localization. To facilitate the disease mechanism investigation, we employed model organism *D. melanogaster,* identified human TREX1 ortholog *cg3165*, and confirmed its vital significance to flies. Then, we expressed human TREX1 and its mutant form TREX1 V235Gfs in flies and used optical coherence microscopy (OCM) to monitor the dynamics of flies’ vascular system. We detected the relapse of fly dorsal vessel, movement impairment, and reduced longevity in TREX1 V235Gfs-expressing transgenic animals. Vascular deterioration and shorter life span recapitulate the RVCL-S manifestations in humans. We have established a robust quantitative *Drosophila* RVCL-S phenotypic system that can potentially serve as a screening platform for drug discovery and drug targets identification.

## Introduction

Retinal vasculopathy with cerebral leukoencephalopathy and systemic manifestations (RVCL-S, RVCL) represents a very special case among rare diseases. It is classified as an ultra-rare disease since it was detected in approximately 30 unrelated families from different countries across the world (1). The disease manifests in highly vascularized tissues including the central nervous system (CNS), retina, liver, kidney (2). The observed symptoms appear between 35 to 50 years of age. Vision impairment, MRI detected brain abnormalities, proteinuria, and liver disease are among the most common symptoms. The severity of the disease correlates with age, leading to premature death (2). Proper diagnosis can be established only based on molecular DNA analyses, which is not a common approach taken by a family physician. Most likely, RVCL-S is under-diagnosed and has much broader distribution among the population (3).

A hereditary syndrome resulting in brain pseudotumors and retinal capillary abnormalities (cerebroretinal vasculopathy, CRV) was first reported and described in 1988 (4). Later, hereditary autosomal dominant vascular retinopathy (HVR), migraine, and Raynaud’s syndrome cases were studied in a large Dutch family (289 family members); the results indicated vascular etiology of this disorder (5). Clinical cases of hereditary endotheliopathy with retinopathy, nephropathy, and stroke (HERNS) were observed in Chinese family (6). HVR, CRV, and HERNS phenotypes were all linked to the same chromosomal region 3p21.1-p21.3 (7). In 2007 these illnesses were designated as retinal vasculopathy with cerebral leukodystrophy (RVCL) and linked to mutations in TREX1 gene resulting in protein C-terminal truncations (8, 9). These mutations are dominant and result in 100% disease penetrance (8). RVCL is not associated with elevated levels of Type I interferons (IFNs) (10). Development of the appropriate therapy remains very challenging, despite the availability of information related to the TREX1 mutations mapping and the protein function. There are no financial incentives for pharmaceutical companies to invest in large scale drug discovery due to the high cost and a small number of RVCL-S patients. One of the directions for RVCL-S treatment development is drug re-purposing, this approach can reduce the time and associated cost. First clinical trial utilizing Aclarubicin, a component of anti-cancer drug cocktails used in China and Japan, to treat RVCL patients started in 2016 (ClinicalTrials.gov, NCT02723448). However, no benefits were observed and the trial did not advance to Phase II (11). Recently, Crizanlizumab, approved for sickle cell anemia treatment, was employed to treat RVCL-S patients in a trial showing a potential to slow the disease progression (12).

Using model organisms to create human disease models has been proven to be an indispensable approach to decipher the disease mechanism on molecular and physiological levels and facilitate the treatment development. Introduction of RVCL-S associated mutation mimicking human TREX1 V235G frameshift into a mouse Trex1 gene resulted in increased mortality and vascular phenotypes in homozygous mice; however, not all pathological features of the disease were detected (13). Studies of TREX1 function on molecular level using cell culture approaches revealed TREX1 mediated DNA damage and subsequent senescence induction caused by a nuclear envelope rupture (14). This may occur naturally in highly mechanically solicited tissues (muscles, etc) due to aging, and also in crowded tissues, *aka* tumors. Chauvin et al. (2024) utilized sophisticated multi-model approach (*Drosophila*, mouse, and cells) to demonstrate the role of RVCL-S causing TREX1protein in accumulation of DNA breaks, cellular senescence induction and loss of specific cell types (11). Regardless of a significant progress in understanding the link between TREX1 mutations and RVCL symptoms, there is still a need for a robust disease model to perform high throughput drug screenings; for example, FDA approved compound libraries assessment could help to prioritize the drug candidates.

*Drosophila melanogaster* (fruit fly) is an invertebrate model organism with well-studied physiology, behavior, sequenced genome, and availability of sophisticated genetic and biochemical tools. Fruit flies possess significant gene conservation with humans and have been successfully used to create human disease models, including rare disease models (15, 16) (17). RVCL-S linked TREX1 gene has an ortholog in *D. melanogaster*, *cg3165,* based on the computational predictions (18). *cg3165* is expressed at all developmental stages, as demonstrated by RNA-Seq data (SI Fig. 1)(19–21). *cg3165* and human TREX1 belong to 3’-5’ exonuclease, DnaQ-like subfamily. Humans have 2 closely related genes, TREX1 and TREX2, their catalytic domains are 40% identical, but TREX2 does not have extended C-terminal region (22). TREX2 has nuclear localization and plays a role in genome stability (23). The C-terminal region of TREX1 contains the transmembrane domain (TM) anchoring it to perinuclear endoplasmic reticulum (24, 25). TREX1 C-terminal truncations lose their cytoplasmic localization and enter the nucleus causing DNA damage and leading to RVCL-S manifestations (11). *cg3165* has significant similarity to TREX1 and TREX2 in N-terminal conserved exonuclease domain, but Alliance of Genome Resources gives higher score to human TREX1 to be an ortholog of *D. melanogaster cg3165* (18). *cg3165* does not have a TM domain (25), likely using different mechanisms to separate nuclear and cytoplasmic activities.

*Drosophila* is the only invertebrate model that has functional and genetically trackable cardiovascular system (26). Flies have open circulatory system with a dorsal vessel spanning along the anterior-posterior axis (27). We focused our research on creating RVCL-S disease model mirroring genetic aberrations and looking for the phenotypic changes in vascular and other relevant organ systems. Robust phenotypes would be used for creating *Drosophila* based RVCL-S screening platform. Previously, we applied optical coherence microscopy (OCM) for non-invasive, *in vivo* visualization of functioning *D. melanogaster* cardiovascular system, producing images with a micron scale resolution (28–32). The imaging data were efficiently processed using deep-learning-based neural network models to create masks of vessel cross section area and extract relevant cardiovascular functional parameters (33–36).

Here, we report the development of an RVCL-S model in *D. melanogaster*. We confirmed the implication of human TREX1 fly ortholog, *cg3165*, in dorsal vessel maintenance, neuromotor regulation, and lifespan prolongation. We have generated a transgenic system, enabling expression of human full length TREX1 and RVCL-S associated C-terminus truncated TREX1 V235G fs in flies. The obtained results have demonstrated the detrimental effects of TREX1 V235G fs on dorsal vessel parameters, neuromotor functions and a lifespan. These instruments can be utilized in search efforts for therapeutic agents to counteract the RVCL-S disease progression.

## Results

### Genetic design of RVCL-S model in *D. melanogaster*

We have identified *Drosophila melanogaster cg3165* as an ortholog of human TREX1 using BLAST and other bioinformatic instruments and searching the genomic and proteomic databases (FlyBase, KEGG, etc.). *cg3165* gene structure is shown in Fig. 1A. The first step in generation of RVCL model was aimed to determine the vital significance of *cg3165* gene for *D. melanogaster*. We performed RNAi mediated CG3165 depletion utilizing UAS/GAL4 genetic technique (37) (Table 1). Ubiquitous depletion (*Act5C> cg3165^RNAi^*) resulted in reduced longevity (SI Fig. 2A) and flies’ impaired locomotor behavior (SI Fig. 2C), consistent only in males. Cardiac vessel specific depletion in *Hand>cg3165^RNAi^* animals did not result in longevity reduction (SI Fig. 2A, B). These initial results, however, suggested the importance of *cg3165* for the flies’ health. Next, we designed a series of transgenic lines with various *D. melanogaster cg3165* and human TREX1 content in order to optimize and establish a robust genetic system representing the RVCL-S disease state. We have generated several genetic combinations, starting from complete removal of fly CG3165 by CRISPR. Though the complete removal of 3’-5’-DNA exonuclease coded by *cg3165* is rather attributable to Aicardi-Goutieres syndrome (38), and is not relevant to the RVCL-S caused by the active enzyme mis-localization, the results obtained with the deletion line were used as a starting point to assess the phenotypic impacts from human TREX1 transgenes. In subsequent steps we gradually added back a copy of *cg3165* gene and introduced transgenic human TREX1 variants. hTREX1 transgenes expression levels were regulated by different GAL4 drivers; details are summarized in Table 1.

**Figure 1.**
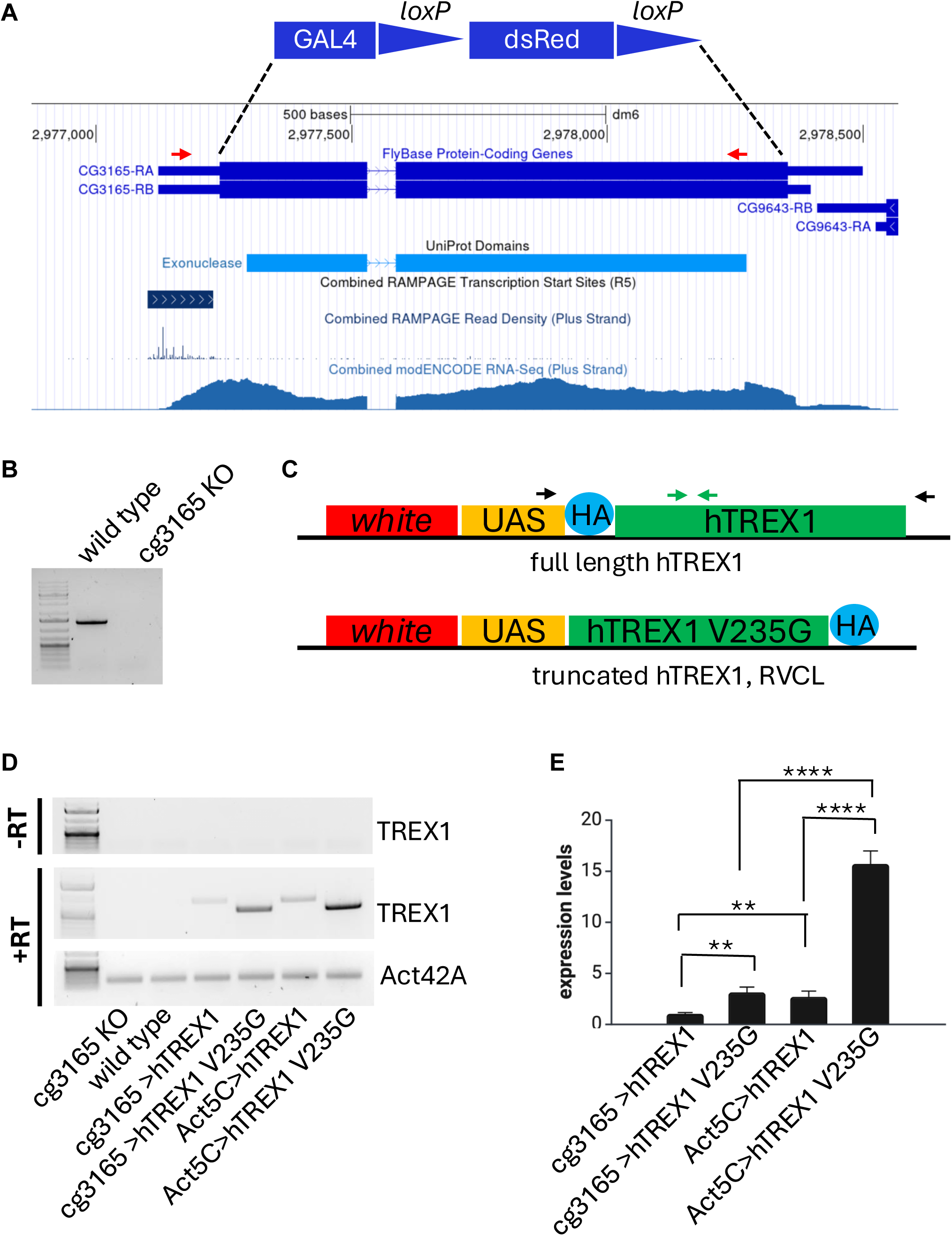
Schematic presentation of generated transgenic constructs and their confirmation after genomic integration and activation. (A) Genomic region containing *cg3165* gene shown in GEP UCSC browser snapshot with selected tracks indicating protein domain, regulatory regions, and RNA-seq coverage. *cg3165* CDS shown as thick blue blocks; it was replaced by GAL4-lox-dsRed-lox sequence shown between dashed lines (functional elements are not in scale). Red arrows indicate PCR primers used to confirm *cg3165* CDS removal. (B) PCR results confirm the removal of CDS fragment. (C) Transgenic constructs used for establishment of UAS-hTREX1 and UAS-hTREX1 V235G fs *Drosophila* stocks. Black arrows indicate RT-PCR primers. Primers used for quantitative RT-PCR are shown in green. (D) RT-PCR results confirm hTREX1 and hTREX1 V235G fs expression. Genotypes are shown below the gel images. *Act42A* gene was used as a reference. Upper panel represents ‘no reverse transcription’ (-RT) control. First lane is the 1 kb Plus DNA Ladder. Primers’ targets are indicated on the right. (E) Quantitative RT-PCR results demonstrate higher expression levels of both TREX1 transgenes when driven by ubiquitous Act5C-GAL4 driver, compared to cg3165-GAL4 driver and show elevated transcription levels of hTREX1 V235G fs compared to hTREX1. Expression levels were normalized to *Act42A*. Bar graphs represent means with SD; ** p< 0.01; **** p< 0.0001.

We have generated a *cg3165 KO-GAL4* strain by knocking out *cg3165* protein coding DNA sequence (CDS) and knocking-in a GAL4 activator sequence. This line plays a dual role: it represents a null *cg3165* mutant (Fig. 1A-B), and, at the same time, it serves as a GAL4 driver regulated by the remaining 5’UTR of *cg3165* gene (Fig. 1A, RAMPAGE evidence track)(39, 40). We utilized this GAL4 driver to activate human TREX1 transgenes in *D. melanogaster* presumably natural spatiotemporal pattern. Human TREX1 transgenes were assembled as following: full length TREX1 or truncated TREX1 G235V fs DNA sequences were placed under the control of UAS element in *Drosophila* transformation vector. Both transgenic constructs were incorporated into *D. melanogaster* genome at *attP2* docking site allowing high expression levels (41) when activated by GAL4 drivers (Fig. 1C, Table 1). *cg3165-GAL4 or Act5C-GAL4* females were crossed to *UAS*-*hTREX1* and *UAS-hTREX1 V235G fs* males and produced viable progeny. We confirmed the expression of *UAS*-*hTREX1* and *UAS-hTREX1 V235G fs* in adult flies (Fig. 1D) and evaluated the RNA levels of both transgenes controlled by cg3165-GAL4 and ubiquitous Act5C-GAL4 (Fig. 1E). As expected, we observed higher transcript levels for Act5C-GAL4 driven transgenes compared to cg3165-GAL4 driven ones. We also noticed strikingly higher expression of RVCL linked *hTREX1 V235G fs* versus normal *hTREX1* under control of both drivers (Fig. 1E). However, we could not detect a truncated hTREX1 V235G fs protein neither with anti-HA (SI Fig. 3A) nor with anti-TREX1(SI Fig. 3B) antibodies by Western blots, where full length hTREX1 produced a band of expected size and intensity (*Act5C> TREX1* is brighter that *cg3165> TREX1*) (SI Fig. 3A-B); these results may reflect the truncated protein instability.

### CG3165 knock-out affects the fly dorsal vessel physiology

In RVCL-S patients with diverse symptoms and manifestations, small blood vessels are affected comprehensively. We set up a goal to characterize the parameters change of the only fruit fly’s vessel in response to the presence of RVCL linked human TREX1. To achieve this, we performed a series of genetic manipulations and assessed our disease model at different building steps. We started with the complete removal of fly CG3165. Young seven day old *cg3165 KO-GAL4* adults’ dorsal vessels were subjected to OCM imaging, males and females were imaged separately (32). Imaging data quantitation suggests that the physiology of the fly dorsal vessel was changed (Fig. 2, SI Video 1-2, SI Fig. 4). We determined several parameters including heart rates (HR), end diastolic area (EDA), end systolic area (ESA), fractional shortening (FS), and arrhythmicity index (AI).

**Figure 2.**
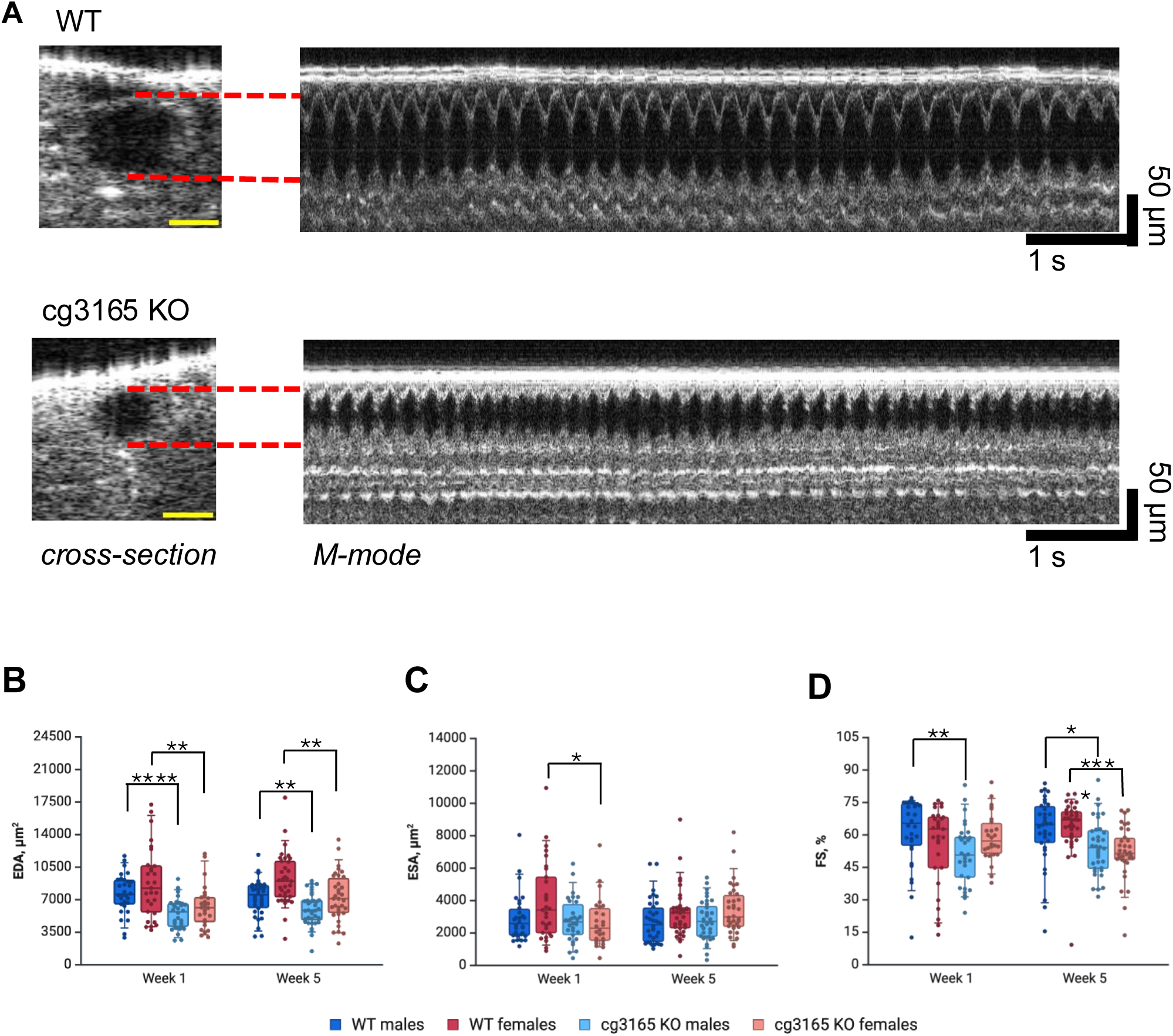
cg3165 knock out affects *D. melanogaster* dorsal vessel functional parameters. (A) Cross-section and M-mode OCM images of WT and cg3165 KO-GAL4 flies (7 days old males shown). Yellow scale bar is 50 um. (B) End diastolic area (EDA) measurements. (C) End systolic area (ESA)measurements. (D) Fractional shortening (FS) parameters. One week and 5 weeks old flies shown in panels B-D; males are shown as blue shade boxes, females shown as red shade boxes. 1 week old sample sizes: WT males n=30, WT females n=29; cg3165 KO-GAL4 WT males n=30, KO-GAL4 WT females n=26. 5 weeks old sample sizes: WT males n=34, WT females n=35; cg3165 KO-GAL4 males n=36, cg3165 KO-GAL4 females n=34. Statistical significance shown as black brackets. * - p< 0.05; ** - p< 0.01; *** - p< 0.001; **** - p< 0.0001.

*cg3165* KO-GAL4 flies demonstrated vessel’s impaired ability to dilate as judged by the EDA measurements; both males and females were affected (Fig. 2B). The average EDA in 1-week-old WT males is 7.7 ± 0.4 x 10^3^ μm^2^ vs 5.5 ± 0.3 x 10^3^ μm^2^ in *cg3165 KO-GAL4* (both n =30, p <0.0001); the females’ average EDA is 8.8 ± 0.7 x 10^3^ μm^2^ vs 6.3 ± 0.5 x 10^3^ μm^2^ respectively (WT n = 29; *cg3165 KO-GAL4* n = 30; p < 0.01). This phenotype was observed in young flies (1-week-old) as well as in aged flies (5-week-old). The average EDA in 5-week-old WT males is 7.2 ± 0.3 x 10^3^ μm^2^ vs 6.0 ± 0.3 x 10^3^ μm^2^ in *cg3165 KO-GAL4* (WT n = 34; *cg3165 KO-GAL4* n = 36; p < 0.01); the females’ average EDA is 9.2 ± 0.5 x 10^3^ μm^2^ vs 7.3 ± 0.4 x 10^3^ μm^2^ respectively (WT n = 35; *cg3165 KO-GAL4* n = 34; p < 0.01).

Reduction of the vessel diameter during maximum contraction, ESA, in *cg3165* KO flies was observed in 1-week-old females (Fig. 2C); WT 1 week old females ESA average is 4.0 ± 0.4 x 10^3^ μm^2^ and *cg3165 KO-GAL4* is 2.7 ± 0.3 x 10^3^ μm^2^ (WT n = 29; *cg3165 KO-GAL4* n = 26; p < 0.05). One week old males have comparable ESA measurements; WT average is 2.9 ± 0.3 x 10^3^ μm^2^ and cg3165 KO-GAL4 is 2.8 ± 0.3 x 10^3^ μm^2^ (WT n = 30; *cg3165 KO-GAL4* n = 30; p > 0.05). Five-week-old males and females ESA were not affected by the removal of CG3165 (Fig. 2C).

Fractional shortening (FS) parameter reflects *Drosophila* dorsal vessel contractility; it measures the percentage difference between the diastolic and systolic states of the fly vessel. We observed impaired contracting ability in 1 week old *cg3165* null males, FS reduction in females was not statistically significant. However, in older flies, both males and females have demonstrated impaired contractility. In 1-week-old WT males, the average FS is 61.5 ± 2.9 % and FS of *cg3165 KO-GAL4* 1-week-old males is significantly lower, 51.0 ± 2.6 % (both genotypes n = 30, p < 0.01). FS of WT and *cg3165 KO-GAL4* 1 week old females is not significantly different, 55.5 ± 3.4 % and 59.0% ± 2.2 % respectively (WT n = 29; *cg3165 KO-GAL4* n = 26; p >0.05, Fig. 2D). The average FS in 5-week-old WT males is 62.0 ± 2.7 % vs 54.3 ± 2.1 % in *cg3165 KO-GAL4* (WT n = 34; *cg3165 KO-GAL4* n = 36; p < 0.05); the females’ average FS is 63.9 ± 2.1% vs 51.9 ± 2.1%, respectively (WT n = 35; *cg3165 KO-GAL4* n = 34; p < 0.0001, Fig. 2D).

Analyzing the other characteristics of fly cardiovascular system, such as heart rates (HR) and arrhythmicity indexes (AI), we detected no significant effects from CG3165 ablation. Of note, the distinct HR were observed between two age groups (1-week-old and 5-week-old flies). In *Drosophila,* HR tends to reduce with age (42, 43); we perceived this in WT and *cg3165* KO-GAL4 flies comparing 1-week-old with 5-week-old males and females, further validating our research tools (SI Fig. 4A). The rhythmicity (AI) displays of *cg3165 KO-GAL4* and WT flies were similar across both age groups (SI Fig. 4B). These results suggest no involvement of *cg3165* in heart regulation *per se*. However, the functioning pattern of the *cg3165* KO-GAL4 flies cardiac vessel is noticeably different from the WT and some phenotypic manifestations (EDA, FS) appear to be related to vessel ‘rigidity’ and are age sensitive. Because the most robust changes were observed in EDA values across both sex and age groups, we decided to use primarily EDA read-out in our following experiments.

### Transgenic hTREX1 rescues the phenotypes caused by *cg3165* knock-out

After we cleared *D.melanogaster* genome of endogenous CG3165 and ascertained the vascular phenotypes, we introduced transgenic human TREX1 or RVCL-S associated TREX1 V235G fs to test the ability of human TREX1 to rescue the vascular phenotypes caused by the *cg3165* knock-out. We crossed *cg3165 KO-GAL4* line carrying 0 copies of *cg3165* with *UAS-hTREX1* or *UAS-hTREX1 V235G fs* line containing hTREX1 transgene and 2 copies of endogenous *cg3165* (SI Fig. 5). Therefore, the resulting *cg3165>TREX1* or *cg3165>TREX1 V235G fs* progeny contains 1 copy of *cg3165* and an activated hTREX1 transgene (Table 1, SI Fig. 5). This balanced genetic design was applied to prevent possible negative effects from an introduction of human protein into fly organism.

To assess the vascular phenotypes, flies were subjected to OCM imaging; *cg3165> hTREX1* (1x *cg3165*, 1x hTREX1) and *cg3165> hTREX1 V235G* (1x *cg3165*, 1x hTREX1 V235G fs) were compared to WT (2x *cg3165*, 0x hTREX1), *cg3165 KO-GAL4* (0x *cg3165,* 0x hTREX1), and *cg3165> yw* (1x *cg3165,* 0x hTREX1). Adult flies’ dorsal vessel was imaged at day 7 (1 week) and day 35 (5 weeks) post eclosure. The EDA measurements are summarized in Fig. 3A-B; SI Video 1-4 contain representative videos. The vessel’s diastolic diameter is increased upon the introduction of a copy of cg3165 and transgenic hTREX1 and TREX1 V235G fs as seen in young males (1-week-old). *cg3165 KO-GAL4* average EDA is 5.5 ± 0.3 x 10^3^ μm^2^ versus 7.0 ± 0.4 x 10^3^ μm^2^ in *cg3165> yw* (both n = 30, p < 0.01), 7.5 ± 0.5 x 10^3^ μm^2^ in *cg3165> hTREX1* (n = 26, p < 0.01), and 7.6 ± 0.5 x 10^3^ μm^2^ in *cg3165> hTREX1 V235G* (n = 25, p <0.01) (Fig. 3A). One-week-old *cg3165> yw*, *cg3165> hTREX1*, and *cg3165> hTREX1 V235G* females also show statistically larger EDA, compared to *cg3165 KO-GAL4* (Fig.3B). However, *cg3165> hTREX1 V235G* average EDA (n = 29) is smaller than *cg3165> yw* (n = 30) hemizygous control (8.2 ± 0.4 x 10^3^ μm^2^ vs 9.9 ± 0.7 x 10^3^ μm^2^, p < 0.05, Fig. 3B). The aged hTREX1 and hTREX1 V235G transgenic flies (5 weeks after eclosion) demonstrate vascular relapse, as the EDA values of *cg3165 >TREX1* and *cg3165 >TREX1 V235G fs* flies are smaller than in control *cg3165 >yw*; this effect is observed for both males and females. The average EDA in 5-week-old *cg3165> yw* males (n = 35) is 8.7 ± 0.4 x 10^3^ μm^2^ versus 7.5 ± 0.4 x 10^3^ μm^2^ in *cg3165> hTREX1* (n = 36), and 7.0 ± 0.4 x 10^3^ μm^2^ in *cg3165> hTREX1 V235G* (n = 35) (p < 0.05 and p < 0.01, respectively); the 5-week-old females’ average EDA in *cg3165> yw* (n = 36) is 11.4 ± 0.5 x 10^3^ μm^2^ versus 8.5 ± 0.4 x 10^3^ μm^2^ in *cg3165> hTREX1* (n = 31), and 9.3 ± 0.5 x 10^3^ μm^2^ in *cg3165> hTREX1 V235G fs* (n = 34, p< 0.0001 and p<0.01, respectively). However, *cg3165> hTREX1* and *cg3165> hTREX1 V235G fs* EDA remains significantly larger than the EDA of 5-week-old *cg3165 KO-GAL4* flies (Fig. 3A-B).

**Figure 3.**
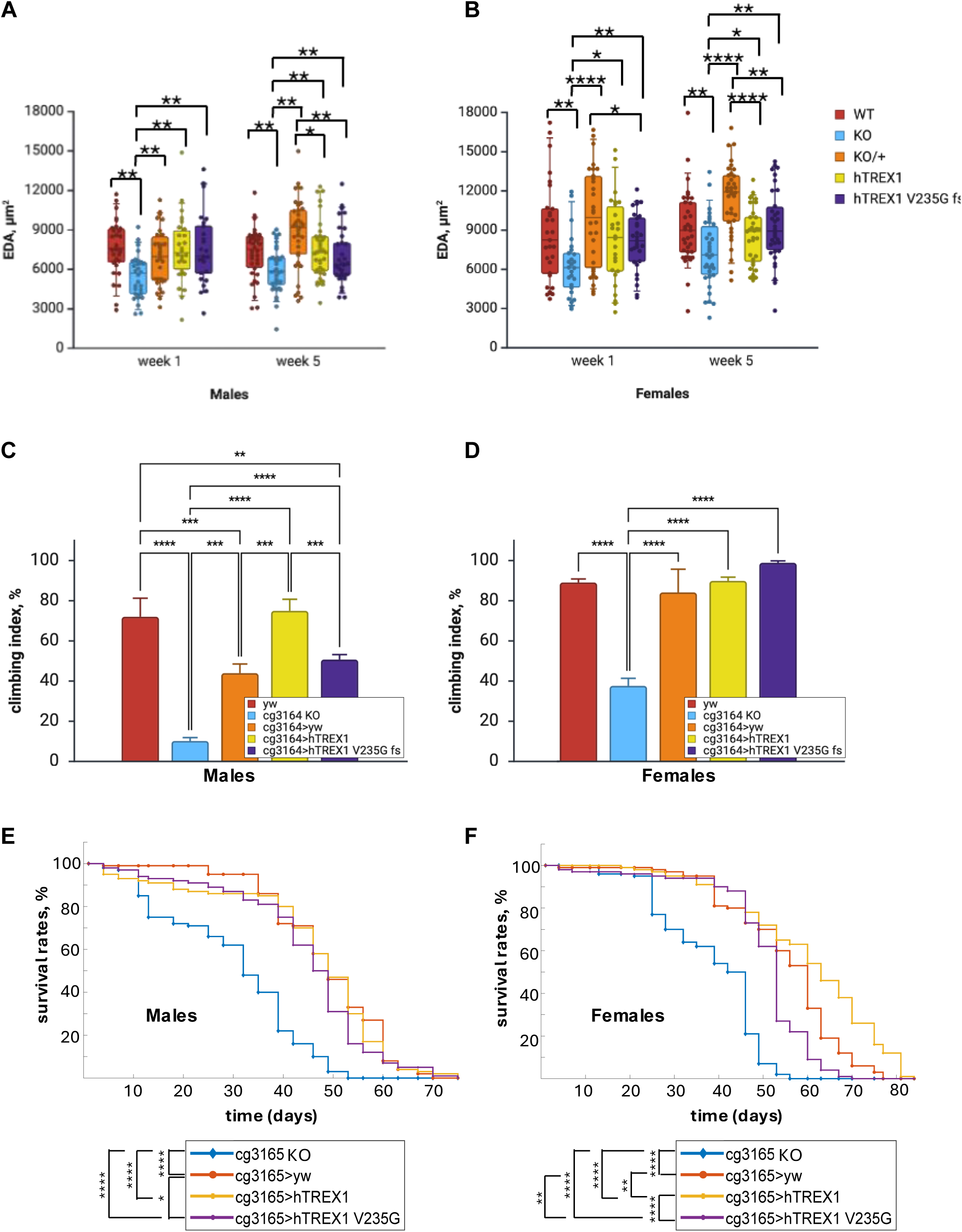
EDA reduction, behavioral impairment, and reduced survival probability observed in CG3165 KO flies are restored by transgenic expression of human TREX1 variants. (A) One-week-old males’ EDA is significantly larger in *cg3165> yw, cg3165> TREX1, and cg3165> TREX1 V235G fs* than in *cg3165 KO*. Five-week-old males carrying hTREX1 and hTREX1 V235G fs demonstrate reduction of EDA compared to *cg3165> yw*. (B) One-week-old females’ EDA is significantly larger in *cg3165> yw, cg3165> TREX1, and cg3165> TREX1 V235G fs* than in *cg3165 KO*. Five-week-old females carrying hTREX1 and hTREX1 V235G fs demonstrate reduction of EDA compared to *cg3165> yw*. WT-*yw*, KO-*cg3165 KO-GAL4*, KO/+ - *cg3165> yw*; *hTREX1 - cg3165> hTREX1*; hTREX1 V235G fs - *cg3165> TREX1 V235G fs*. 1 week old sample sizes: WT males n=30, KO males n=30, KO/+ males n=30, hTREX1 males n=26, hTREX1 V235G fs males n=25; WT females n=29, KO females n=26, KO/+ females n=30, hTREX1 females n=26, hTREX1 V235G fs females n=29. 5 weeks old sample sizes: WT males n=34, KO males n=36, KO/+ males n=35, hTREX1 males n=36, hTREX1 V235G fs males n=34; WT females n=35, KO females n=34, KO/+ females n=36, hTREX1 females n=31, hTREX1 V235G fs females n=34. (C) *cg3165* knock-out males show impaired locomotor behavior compared to WT (*yw*); hemizygous *cg3165* (*cg3165> yw*) males demonstrate significant improvement. hTREX1 introduction (*cg3165>* hTREX1) results in climbing ability improvement comparable to WT levels. Males carrying RVCL-S associated hTREX1 V235G fs (*cg3165> TREX1 V235G fs*) demonstrate partial climbing ability restoration. (D) *cg3165* knock-out females show impaired climbing compared to WT (*yw*). Normal locomotor behavior is observed in *cg3165> yw, cg3165> TREX1, and cg3165> TREX1 V235G fs* animals. Bar graphs (A-B) show means with SD. (E) Kaplan-Meier survival curves of *cg3165* KO, *cg3165 >yw*, *cg3165 >hTREX1* and *cg3165 > hTREX1 V235G fs* males. Survival rates of *cg3165* knock-out males are significantly reduced. Upon introduction of 1 copy of *cg3165* (*cg3165 >yw*) and transgenic hTREX1 (*cg3165 >hTREX1*), or hTREX1 V235G fs (*cg3165 > hTREX1 V235G fs*) the survival probability is increased. (F) Kaplan-Meier survival curves of *cg3165* KO, *cg3165 >yw*, *cg3165 >hTREX1* and *cg3165 > hTREX1 V235G fs* females. Survival rates of *cg3165* knock-out females are significantly reduced. Upon introduction of 1 copy of *cg3165* (*cg3165 >yw*) and transgenic hTREX1 (*cg3165 >hTREX1*), or hTREX1 V235G fs (*cg3165 > hTREX1 V235G fs*) the survival probability is increased. hTREX1 introduction has the strongest impact on longevity; hTREX1 V235G fs effect is the weakest. Statistical significance shown as black brackets.: * - p< 0.05; ** - p< 0.01; *** - p< 0.001; **** - p< 0.0001.

Analyzing the EDA phenotypes in current hTREX1 expression system, we observed the rescue by the hTREX1 and hTREX1 V235G fs of a vascular phenotype caused by CG3165 knock-out, though we did not detect the significant differences between the full length hTREX1 and hTREX1 V235G fs efficiency. Comparing 1-week and 5-week post eclosure groups we noticed aging-related detrimental effects caused by hTREX1 V235G fs, and also by a full length hTREX1 in males and females (Fig. 3A-B).

RVCL affects multiple organ systems and has neurological manifestations including brain dysfunction (9). Fruit flies’ survival greatly depends on their motor functions; this is achieved by tight coordination between the CNS processing external signals and sending instructions, motor neurons, and muscles performing certain actions. Disruption of these processes results in motor defects (44). We performed a climbing assay to test the ability of TREX1 mutant to maintain a normal behavioral pattern of negative geotaxis. We observed that the *cg3165* knock-out resulted in significantly impaired climbing ability of *cg3165 KO-GAL4* flies; climbing index dropped from WT 72.0 ± 9.2% to 10.1 ± 1.7% (p < 0.0001) in *cg3165 KO-GAL4* males; from 89.1 ± 1.8% to 37.6 ± 3.7% (p < 0.0001) in females (Fig. 3C-D), suggesting that the *cg3165* gene plays a broad role affecting the neurological networks and/or muscle tissue condition. The climbing ability has significantly improved in hemizygous *cg3165 >yw* (climbing index is 44.0 ± 4.5% for males (p < 0.001), 84.1 ± 11.5% (p < 0.0001) for females) and in flies expressing both hTREX1 variants: *cg3165 >hTREX1* (75.0 ± 5.7%, p < 0.0001 for males; 89.9 ± 1.8%, p < 0.0001 for females), *cg3165 >hTREX1 V235G fs* flies (50.7 ± 2.5% (p < 0.0001) for males; 98.9 ± 1.0% (p < 0.0001) for females) (Fig. 3C-D). In particular, males show remarkable gene dose sensitivity in restoration of locomotor behavioral patterns. Adding hTREX1 transgenic copy significantly increased the climbing index compared to *cg3165 KO-GAL4 and cg3165 >yw,* while the effect from RVCL-S associated hTREX1 V235G fs was significantly less compared to hTREX1 carrying flies (Fig. 3C).

Lifespan is a robust indicator of aging rates in fly population. We performed the longevity study (25C, 70 % humidity) to determine the effects of *cg3165* knock-out and introduction of human transgenic copies of full length or truncated hTREX1 forms. Our initial experiments implied that CG3165 depletion reduced the lifespan, but only in males (SI Fig. 2A). Complete CG3165 removal significantly reduces the life span of *cg3165 KO-GAL4* flies (Fig. 3E-F, SI Table 1), males and females, compared to *cg3165 >yw* (p < 0.0001 for males, p < 0.0001 for females)*, cg3165 >hTREX1* (p < 0.0001 for males, p < 0.0001 for females) and *cg3165 > hTREX1 V235G fs* flies (p < 0.0001 for males, p < 0.0001 for females). Survival probability was increased in *cg3165 >hTREX1* females, compared to *cg3165 >yw* control (p < 0.01, Fig. 3F, SI Table 1); females carrying RVCL associated hTREX1 V235G fs demonstrate reduced longevity compared to *cg3165 >yw* (p < 0.01), and to *cg3165 >hTREX1* (p < 0.0001). *cg3165 >yw, cg3165 KO-GAL4>hTREX1* and *cg3165 >hTREX1 V235G fs* males show similar lifespan patterns (Fig. 3E), close to average *Drosophila melanogaster* male lifespan at 25C (45).

### Over-expression of human TREX1 transgenes show the age dependent phenotypes

To enhance the phenotypic effects from the ectopic hTREX1 and hTREX1 V235G fs expression we decided to use strong ubiquitous Act5C-GAL4 driver to increase the transgene transcription rates (Fig. 1E). We performed crosses between *Act5C-GAL4* females and *UAS-hTREX1* and *UAS-hTREX1 V235G fs* males (Table 1, SI Fig.6). One needs to point out, that these genetic configurations include 2 copies of *cg3165* (SI Fig. 6), not 1, as in *cg3165 >hTREX1 and cg3165 >hTREX1 V235G fs* flies (Table 1, SI Fig.5). Higher *Act5C-GAL4 driven UAS-hTREX1* and *UAS-hTREX1 V235G fs* expression levels compared to *cg3165 >hTREX1 and cg3165 >hTREX1 V235G fs* flies’ levels (Fig. 1E) did not lead to an increased lethality during early developmental stages of hTREX1 and hTREX1 V235G fs expressing progeny compared to non-transgenic siblings. Obtained *Act5C >hTREX1 V235G fs* flies’ longevity measurements have shown notably shorter lifespan relative to *Act5C>hTREX1* (p < 0.0001 for males and p < 0.0001 for females), and *Act5C>yw* animals (p < 0.0001 for males and p < 0.001 for females) (Fig. 4A-B, SI Table 2). hTREX1 expressing females have demonstrated increased longevity compared to control *Act5C>yw* (p < 0.001) (Fig. 4B), while *Act5C >hTREX1* males’ lifespan was similar to the control *Act5C>yw* males (p > 0.05) (Fig. 4A). These results clearly demonstrate the negative effects of RVCL-S linked hTREX1 V235G fs on flies’ survival probability.

**Figure 4.**
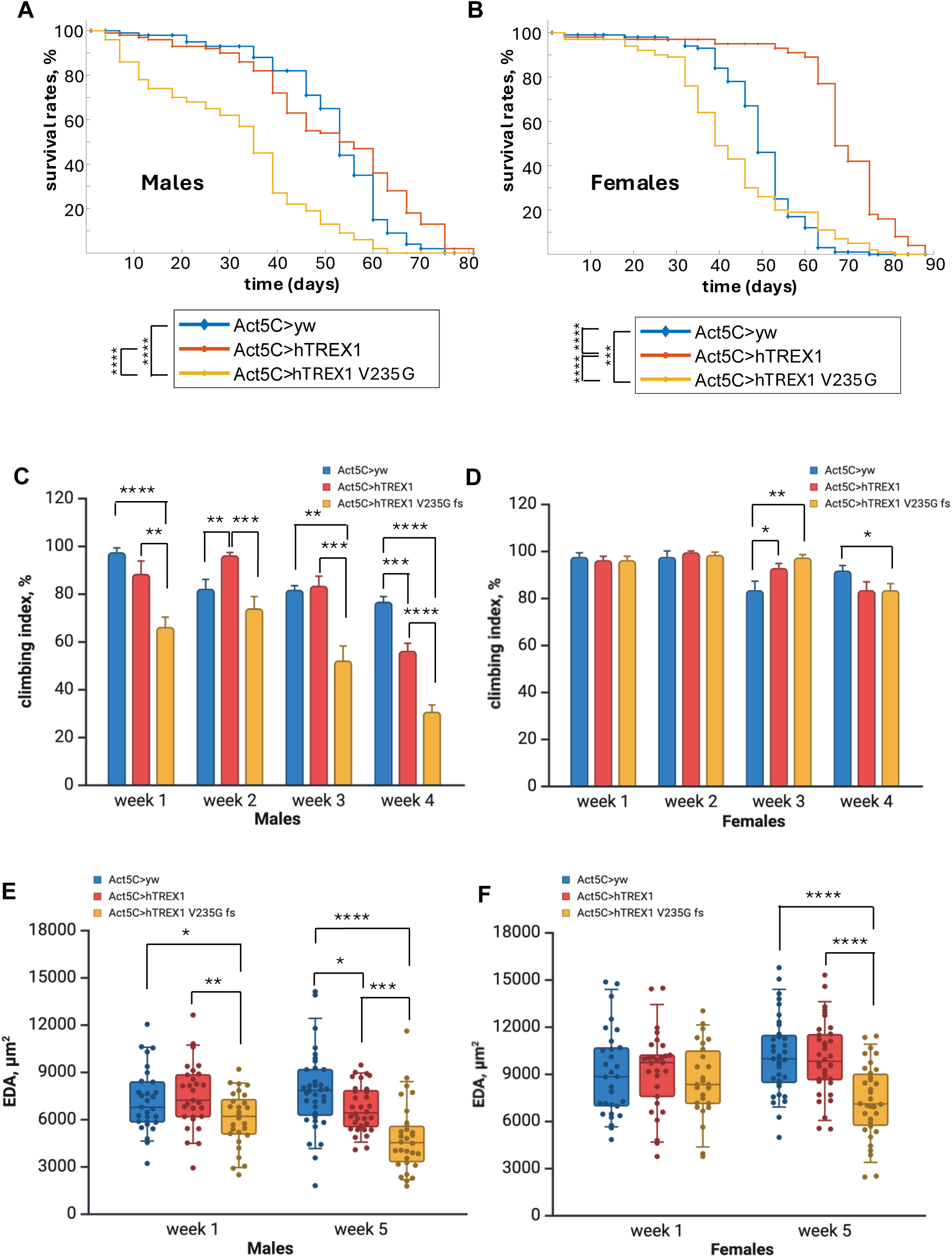
Over-expression of RVCL linked hTREX1 V235G fs reduces survival probability, movement impairment, and decrease of dorsal vessel EDA. In aged flies, over-expression of ‘healthy’ hTREX1 transgene negatively affects climbing ability and results in smaller EDA. (A) Kaplan-Meier survival curves of *Act5C>yw* (blue line), *Act5C>hTREX1* (orange line) and *Act5C >hTREX1 V235G fs* (yellow line) males. (B) Kaplan-Meier survival curves of *Act5C>yw* (blue line), *Act5C>hTREX1* (orange line) and *Act5C >hTREX1 V235G fs* (yellow line) females. C- Climbing assay results in *Act5C*>*yw*, *Act5C*>*hTREX1*, *Act5C* >*hTREX1 V235G fs* males. D- Climbing assay results in *Act5C*>*yw*, *Act5C*>*hTREX1*, *Act5C* >*hTREX1 V235G fs* females. Bar graphs (C, D) represent means with SEM. (D) End Diastolic Area (EDA) measurements of *Act5C*>*yw*, *Act5C*>*hTREX1*, *Act5C* >*hTREX1 V235G fs* males. (E) End Diastolic Area (EDA) measurements of *Act5C*>*yw*, *Act5C*>*hTREX1*, *Act5C* >*hTREX1 V235G fs* females. 1 week old groups sample sizes: *Act5C*>*yw* males n=29, *Act5C*>*hTREX1* males n=29, *Act5C* >*hTREX1 V235G fs* males n=28; WT females n=30, hTREX1 females n=29, *Act5C* >*hTREX1 V235G fs* females n=26. 5 week old groups sample sizes: *Act5C*>*yw* males n=25, *Act5C*>*hTREX1* males n=32, *Act5C* >*hTREX1 V235G fs* males n=30; *Act5C*>*yw* n=35, *Act5C*>*hTREX1* females n=36, *Act5C* >*hTREX1 V235G fs* females n=34. Statistical significance shown as black brackets. * - p< 0.05; ** - p< 0.01; *** - p< 0.001; **** - p< 0.0001.

Behavioral tests were performed on aging flies, from week 1 to week 4 after eclosion, to assess the effect of mutant hTREX1 V235G fs over-expression (Fig. 4C-D). Impaired climbing ability was observed in *Act5C >hTREX1 V235G fs* males starting from the young age, at week 1 and persisted to the end point. *Act5C >hTREX1 V235G fs* females started showing the moving impairment much later, at week 4. The *Act5C>hTREX1* and *Act5C >hTREX1 V235G fs* flies’ responses to climbing challenge were similar to the results observed for 1 week old *cg3165 >hTREX1* and *cg3165 >hTREX1 V235G fs* flies (Fig. 3C-D). Males were more sensitive to the detrimental effect of RVCL-S linked hTREX1 mutation. We also noticed the age correlated negative impact from hTREX1 over-expression in both males and females, at week 4 after eclosion, compared to *Act5C>yw* control (Fig. 4C).

Considering the physiological impacts of hTREX1 isoforms over-expression, we performed the OCM imaging of flies’ cardiovascular system at week 1 and week 5 after eclosion. The EDA measurements extracted from the processed imaging data are summarized in Fig. 4E-F, SI Video 5-7. At week 1, we observed the EDA reduction in *Act5C >hTREX1 V235G fs* (RVCL) males (6.0 ± 0.3 x 10^3^ μm^2^, n = 28) relatively to control *Act5C>yw* (7.2 ± 0.4 x 10^3^ μm^2^, n = 29, p < 0.05) and *Act5C>hTREX1* (7.5 ± 0.4 x 10^3^ μm^2^, n = 29, p < 0.005) (Fig. 4E); females average EDA were: *Act5C>yw* - 9.0±0.5 x 10^3^ μm^2^ (n = 30), *Act5C>hTREX1 -* 9.2 ± 0.5 x 10^3^ μm^2^ (n = 29), *Act5C>hTREX1 V235G fs -* (8.6 ± 0.5 x 10^3^ μm^2^, n = 26); the EDA reduction in *Act5C>hTREX1 V235G fs* females is not statistically significant (p>0.05) (Fig. 4F). At week 5, we continued to see EDA reduction in males carrying hTREX1 V235G fs (RVCL). The average EDA of *Act5C>hTREX1 V235G fs* 5-week-old males are 4.8± 0.4 x 10^3^ μm^2^ (n = 30). It is significantly smaller than the average EDA of 5-week-old *Act5C>yw* males (7.8 ± 0.4 x 10^3^ μm^2^, n = 35, p < 0.0001), and the average EDA of 5-week-old *Act5C>hTREX1* males (6.7 ± 0.3 x 10^3^ μm^2^, n = 32, p < 0.001). We also observed the negative effects on EDA in aged *Act5C>hTREX1* males, expressing ‘healthy’ hTREX1. The average EDA of 5-week-old *Act5C>hTREX1* males (6.7 ± 0.3 x 10^3^ μm^2^, n = 32) is smaller than the average EDA of 5-week-old *Act5C>yw* males, (7.8 ± 0.4 x 10^3^ μm^2^, n = 35, p < 0.05, Fig. 4E). The detrimental effect of hTREX1 V235G fs expression became evident in aged females. The average EDA of *Act5C>hTREX1 V235G fs* 5-week-old females are 7.3 ± 0.4 x 10^3^ μm^2^ (n= 34). It is significantly smaller than the average EDA of 5-week-old *Act5C>yw* females (10.2 ± 0.4 x 10^3^ μm^2^, n = 35, p < 0.0001), and the average EDA of 5-week-old *Act5C>hTREX1* females (10.0 ± 0.4 x 10^3^ μm^2^, n = 36, p < 0.0001) (Fig. 4F).

Overall, the longevity test, behavioral assays, and OCM imaging results point to the hTREX1 V235G fs as an adverse modulator of physiological processes in adult flies.

## Discussion

We took advantage of *Drosophila melanogaster* genetic capability and an innovative imaging technology, OCM, to create an RVCL-S disease model that will facilitate the screening for therapeutics and could be used to study the disease progression. We have identified *D. melanogaster* ortholog of human TREX1 gene linked to the disorder, *cg3165,* and confirmed its significance for the flies’ vital functions. The removal of *cg3165* CDS lead to fly’s cardiovascular system changes comprehensively evaluated by OCM. Dorsal vessel parameter deviations were used as a phenotypic read-out for the RVCL-S fly model building, where the end diastolic area (EDA) proved to be the most robust.

*Drosophila* transgenic lines, carrying the full length human TREX1 and RVCL-S associated truncated TREX 1 V235G fs were generated. We focused our efforts on uncovering the distinct phenotypes caused by the full length and the mutant hTREX1. We detected that the flies’ vascular integrity was restored upon the introduction of both human TREX1 transgenes and/or a copy of *cg3165* (Fig. 3, SI Fig. 5) compared to *cg3165* null. However, we could not distinguish between the impacts from the hTREX1 and hTREX1 V235G fs at this point, the EDA parameters of *cg3165> hTREX1* and *cg3165> hTREX1 V235G fs* animals were similar.

We extended our assays to behavioral tests based on the rational that RVCL-S affects cerebral functions (1). Upon *cg3165* knock-out, flies have demonstrated a significant movement impairment at week 1 after eclosure compared to the WT (Fig. 3C-D). Adding a copy of *cg3165*, hTREX1, or hTREX1 V235G fs improved the climbing ability compared to the *cg3165* KO. In males, we observed gradual effects: *cg3165 >yw* demonstrated some improvement; hTREX1 carrying flies’ climbing index was the highest; but hTREX1 V235G fs flies’ climbing ability was significantly lower than of hTREX1 carrying animals. These results suggested strong positive correlation between *Drosophila* neuromotor regulation and transgenic hTREX1 presence and a negative impact from hTREX1 V235G.

In addition to the experiments scrutinizing the organ specific functions related to the RVCL-S, we performed a longevity study of flies with various hTREX1 genetic content (Fig. 3E-F). *cg3165* KO animals are homozygous viable, but their lifespan is significantly shorter. Adding back *cg3165*, hTREX1, or hTREX1 V235G fs prolongs the lifespan. The impact is higher in *cg3165> hTREX1* and is less in *cg3165> hTREX1 V235G fs* flies relatively to the *cg3165>yw* control as seen in females. Longevity is not a phenotype that can be used for screening purposes, but it is very reliable method to strengthen the RVCL-S model in *Drosophila.* RVCL-S patients have decreased life expectancy (2), and our results with hTREX1 V235G fs carrying flies reflect this disease aspect.

Following the observed trends in our results, we changed the genetic content of the experimental setup; instead of maintaining the gene expression pattern and copy numbers (1 copy of endogenous *cg3165* and a human hTREX1/ hTREX1 V235G fs transgene controlled by 5’UTR cg3165), we utilized ubiquitous *Act5C-GAL4* driver to over-express human TREX1 constructs (SI Fig. 6) anticipating to obtain more robust phenotypes. The longevity evaluation of *Act5C>yw* (control), *Act5C> hTREX1*, and *Act5C> hTREX1 V235G fs* flies re-confirmed the negative impact of hTREX1 V235G fs expression on *Drosophila* survival rates (Fig. 4A-B). The locomotor behavior tests performed during week 1 through week 4 demonstrated progressive impairment of climbing ability of hTREX1 V235G carrying males relatively to hTREX1 carrying and control animals (Fig. 4C).

The OCM imaging results of *Act5C>yw*, *Act5C> hTREX1*, and *Act5C> hTREX1 V235G fs* clearly demonstrated the detrimental effect of hTREX1 V235G fs on dorsal vessel dilation in 1-week-old males compared to hTREX1 and ‘no transgene’ control (Fig. 4E). Considering the late onset of RVCL-S manifestations in humans, in aged (5-week-old) flies, we detected the TREX1 V235G fs facilitated the EDA reduction in both males and females (Fig.4E-F).

To build an RVCL-S research model, we devised two *Drosophila* UAS/GAL4 expression systems of human TREX1 and TREX1 V235G fs proteins: *cg3165* and *Act5C* promoter based. The Act5C over-expression approach proved to be more efficient to detect the detrimental effect of RVCL-S linked hTREX1 V235G fs on vascular phenotypes. But the more robust EDA phenotype appearance could also be affected by the fly *cg3165* gene copy numbers in the background (*i.e.*, amount of 3’-5’-DNA exonuclease). McGlasson *et al.* (2025) have recently shown that mono-allelic truncating mutations in TREX1 require intact nuclease activity in order to cause endothelial disease (46). The EDA measurements, shown in Fig. 3A-B, do not indicate any differences between *cg3165> hTREX1* and *cg3165> hTREX1 V235G fs* animals in any sex or age groups. Meanwhile, the dorsal vessel deterioration (smaller EDA) in *Act5C> hTREX1 V235G fs* is obvious in young and aged males, and aged females, compared to of *Act5C> hTREX1* (Fig. 4E-F). The absence of EDA phenotypic differences in the 1^st^ case correlates with *cg3165* haplodeficiency (SI Fig. 5) and, therefore, lower level the exonuclease activity. The *Act5C* based expression system includes two copies of *cg3165* (SI Fig. 6) and has higher levels 3’-5’-DNA exonuclease most likely contributing to the stronger effects in *Act5C> hTREX1 V235G fs* flies.

RVCL-S manifestations increase with aging leading to premature death (2). Comparing two *Drosophila* age groups (1 week and 5 weeks after eclosure), we noticed the vessel deterioration in older flies carrying full length hTREX1 compared to the ‘no transgene’ controls (Fig.3A-B and Fig. 4E). Similar observations were made by Chauvin et al. (2024) (11) when hTREX1 was expressed in *D. melanogaster* eye tissue and caused a rough phenotype, though less severe that of RVCL-S isoform, but distinct from the normal state. The authors observed hTREX1 nuclear mis localization, however less pronounced than in hTREX1 V235G fs containing cells. The vascular damage in aged hTREX1 carrying flies might occur as a consequence of hTREX1 mis localization to the nucleus by the mechanisms described in (14) due to significant mechanical stress within the vessel tissues.

In summary, we have created an experimental methodology aimed to facilitate the development of a treatment for a rare genetic disorder, RVCL-S. Current work represents an interdisciplinary approach, where humans’ medical problems are addressed using *Drosophila* model organism through the methods of genetics, bioinformatics, biophysics and others. Through optimization of these tools, we have built a model that in the future could be used for testing chemical compounds. Our system also allows us to conduct further research on molecular level to identify the druggable gene targets.

## Materials and methods

### Drosophila stocks

Fly stocks were maintained on standard cornmeal media at room temperature. *y[1] w[*]; P{Act5C-GAL4-w}E1/CyO* driver was obtained from Bloomington Drosophila Stock Center, stock #25374. *cg3165* CDS deletion and GAL4 knock-in was achieved by CRISPR and performed by WellGenetics (Taiwan) resulting in *w^1118^; cg3165^KO^ GAL4 loxP RFP loxP/ CyO* stock. To create transgenic stocks carrying human TREX1 gene, we used a plasmid pcDNA3.1 N-HA-human TREX1 (WT) provided by Jonathan Miner (University of Pennsylvania Perelman School of Medicine) as a source. We performed seamless cloning using NEBuilder® HiFi DNA Assembly Master Mix (NEB, E2621S). HA-hTREX1 DNA fragment was amplified with 5’-GGG AAT TGG GAA TTC GTT AAC ACT AGC GTT TAA ACT TAA GCT TGC CAC CAT GTA CCC-3’ and 5’-ATC CTC TAG AGG TAC CCG CGG CCG CCA CTG TGC T-3’ primers and assembled with pUAST-attB (#1419, DGRC) cut with *Xho*I, *Bgl*II. To create transgenic stocks carrying human TREX1 G235V fs gene we used a plasmid pcDNA3.1 hTREX1 V235Gfs C-HA provided by Jonathan Miner. hTREX1 V235Gfs C-HA DNA fragment was amplified with 5’-TAG GGA ATT GGG AAT TCG TTA ACA CTA GCC ACC ATG GG-3’ and 5’-ATC CTC TAG AGG TAC CCA GCG GGT TTA TCA AGC GTA AT-3’ and assembled with pUAST-attB/ *Xho*I, *Bgl*II. Resulting *Drosophila* transformation vectors were microinjected into fly embryos by BestGene Inc. Phi31 mediated cassette exchange occurred at attP2 landing pad. Transgenes integration was verified by PCR and sequencing. *yw; UAS-HA-hTREX1 w^+^ y^+t7.7^ attP2/TM6B* and *yw; UAS-hTREX1 V325G fs w^+^ y^+t7.7^ attP2/TM6B* stocks were established.

### Genetic crosses

*w^1118^; cg3165^KO^ GAL4 loxP RFP loxP/ CyO* females were crossed to *yw; UAS-HA-hTREX1 w^+^ y^+t7.7^ attP2/TM6B* or *yw; UAS-hTREX1 V325G fs w^+^ y^+t7.7^ attP2/TM6B.* Non-*Cy*, non-*Tb* progeny were collected for the experiments. Non-*Cy* progeny from *w^1118^; cg3165^KO^ GAL4 loxP RFP loxP/ CyO* and *yw* cross served as the genetic control.

*yw; P{Act5C-GAL4-w}E1/CyO* females were crossed to *yw; UAS-HA-hTREX1 w^+^ y^+t7.7^ attP2/TM6B Tb.* Non-*Cy*, non-*Tb* progeny was collected for imaging and other experiments. Same crossing scheme was applied for hTREX1 G235V fs carrying flies. Non-*Cy* progeny from *yw; P{Act5C-GAL4-w}E1/CyO* and *yw* cross served as the genetic control.

### OCM imaging

Adult flies were imaged using our custom-built SD-OCM system. Broadband light was sent from a super luminescent diode (SLD) with a center wavelength of 850 nm and a bandwidth of 165 nm (Superlum, cBLMD-T-850-HP). A 10x objective focused light onto the sample stage. Light interference from sample and reference arms was measured using a spectrometer and 2048-pixel line-scan camera (Wasatch Photonics, CS800-840/180-80-OC2K-U3). The lateral resolution was measured as ∼2.8 µm and the axial resolution was ∼3.3 µm in tissue. Measured system sensitivity was ∼95.1 dB.

Flies were fixed to a glass slide by attaching their wings to the slide with rubber cement (Elmer’s, Rubber Cement). The OCM beam was positioned over the A1 segment of the heart. M-Mode imaging was performed using 128 A scans and 2000 B scans with an exposure time of 50 µs and a frame rate of 125 frames per second, for an approximately 16-second-long recording per collection.

### OCM image processing

Raw OCM data was processed into images using custom lab MATLAB code. To quantify relevant heart parameters, the heart area was segmented in each image using FlyNet3.0 to create masks of the heart area. Once masks were created, they were resized, such that each pixel was equivalent to 1 micron. Heart area over time was plotted, and peak (maximum area) and valley (minimum area) points were identified. Heart rate was calculated using the inverse of the distance between valley areas. End diastolic area (EDA) was calculated as the average area in µm^3^ at each peak, and end systolic area (ESA) was calculated as the average area in µm^3^ at each valley. Fractional shortening (FS) was calculated using the following equation:

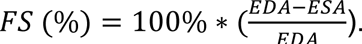

### PCR

hTREX1 transgenes were verified by PCR using GoTaq Master Mix (Promega, M7122) and primers 5’-CCT GCA GGTCGGAGT ACT GT-3’ and 5’-GGA AAG TCC TTG GGG TCT TC-3’ following manufacturer’s instructions.

### RT-PCR

Total RNA was extracted from adult flies using TRI Reagent (Sigma-Aldrich, T9424-25ML) according to manufacturer’s instructions. cDNA synthesis was performed using QuantiTect Reverse Transcription Kit (Qiagen, 205311) according to the manual. Gene specific PCRs were done using GoTaq Master Mix and primers 5’-AGC GAG ATC ACA GGT CTG AG-3’ and 5’-ACC ACT GCT CCC AT CAT CA-3’ to detect hTREX1 transgenes, and 5’-TGC CCA TTT ATG AGG GCT AC-5’ and 5’-ATC TCC TGC TCG AAG TCC AA-3’ specific for Actin 42A gene serving as a control. qRT-PCR was performed on StepOnePlus System (Applied Biosystems) using QuantiTect SYBR Green PCR Kit (Qiagen, 204143) and hTREX1 specific primers, 5’-GCATGGGCGTCAATGTTTTG-3’ and 5’-TGCTATCCACACAGAAGGCA-3’. Actin 42A gene served as reference.

### Western blots

Total protein extracts were obtained from adult flies as follows: flies were homogenized in 1X Laemmli Sample Buffer (Bio-Rad, 1610737) and boiled for 5 min. Proteins were resolved by size in 10% SDS-PAGE and transferred to nitrocellulose membrane (Bio-Rad, 1620112). Membranes we blocked in 5% Blotto (Santa Cruz Biotechnology, sc-2324) in TBS-T. Anti-HA rMs-IgG1-s (DSHB) antibodies were used at 1:1,000 dilution; anti-LaminC (LC28.26-s, DSHB) at 1:1,000; anti-TREX1 (D8E2O) Rabbit mAb #15107 (Cell Signaling Technology) in 2.5% Blotto. Secondary Peroxidase-conjugated AffiniPure Goat Anti-mouse IgG (Jackson ImmunoResearch, 115-035-003) or Peroxidase-conjugated AffiniPure Goat Anti-rabbit IgG (Jackson ImmunoResearch, 111-035-003) were diluted to 1:100,000. SuperSignal™ West Femto Maximum Sensitivity Substrate (Thermo Scientific, 34095) was used for signal development. Signal visualization was performed on iBright750 imaging system.

### Climbing assay

20 flies were placed in an empty vial, left to recover from CO_2_ anesthesia for ∼15 min, gently banged down to bring the flies to the bottom of the vial and then flies were let to climb up the wall. Short videos were recorded and used to calculate the percentage of flies crossed the horizontal line drawn at 2.5 cm height at 10 sec. Multiple vials were taped together. Males and females were tested separately. Bar graphs were created in https://BioRender.com.

### Longevity assay

100 males and 100 females eclosed within 24h were placed in fresh vials, 33-34 flies per vial and kept at 25C 70% humidity. Flies transfer was done every 3-4 days; numbers of dead flies were recorded.

### Statistical analyses

For climbing assay and heart function analyses, two-sample student’s t tests were performed with a 95% confidence level. For longevity assay, the age of flies was tracked for each dataset, and we calculated the Kaplan-Meier survival curve, and performed a log-rank test and two-sample two-tailed *t*-test, also with a 95% confidence level.

## Funding

This research was supported by National Institutes of Health grant R01-HL156265 (C. Z.); National Institutes of Health grant R01AI143982 (J.J.M.); National Institutes of Health grant R01NS131480 (J.J.M.); Clayco Foundation Innovative Research Award (C. Z.); gift from the Clayco Foundation (J.J.M.); National Science Foundation Graduate Research Fellowship Program (A. Matt); Washington University in St. Louis startup fund (C. Z.); Penn Colton Center for Autoimmunity pilot award (J.J.M.); Penn RVCL Sisters Fund (J.J.M.).

We thank Joanna Chen and Dante Zou for their help with the experiments. We are very grateful to the organizers and participants of the first International RVCL-S Meeting (Leiden, Netherlands, 2024) for helpful and enriching discussions.

## Data availability

Data underlying the results presented in this paper are not publicly available at this time but may be obtained from the authors upon reasonable request.

---

Table 1. Genetic configuration UAS/GAL4 system for RNAi mediated depletion of *cg3165* and expression of human TREX1 transgenes in *D. melanogaster* Females from GAL4 driver lines listed in 1^st^ column were crossed to males shown in the upper row. The resulting copy numbers of fly *cg3165* and human TREX1 content of the progeny is described.

